# Early replacement of West Eurasian male Y chromosomes from the east

**DOI:** 10.1101/867317

**Authors:** Pille Hallast, Anastasia Agdzhoyan, Oleg Balanovsky, Yali Xue, Chris Tyler-Smith

## Abstract

The genomes of humans outside Africa originated almost entirely from a single migration out ∼50,000-60,000 years ago^1,2^, followed closely by mixture with Neanderthals contributing ∼2% to all non-Africans^3,4^. However, the details of this initial migration remain poorly-understood because no ancient DNA analyses are available from this key time period, and present-day autosomal data are uninformative due to subsequent population movements/reshaping^5^. One locus, however, does retain extensive information from this early period: the Y-chromosome, where a detailed calibrated phylogeny has been constructed^6^. Three present-day Y lineages were carried by the initial migration: the rare haplogroup D, the moderately rare C, and the very common FT lineage which now dominates most non-African populations^6,7^. We show that phylogenetic analyses of haplogroup C, D and FT sequences, including very rare deep-rooting lineages, together with phylogeographic analyses of ancient and present-day non-African Y-chromosomes, all point to East/South-east Asia as the origin 50,000-55,000 years ago of all known non-African male lineages (apart from recent migrants). This implies that the initial Y lineages in populations between Africa and eastern Asia have been entirely replaced by lineages from the east, contrasting with the expectations of the serial-founder model^8,9^, and thus informing and constraining models of the initial expansion.

## Main

A consensus view has emerged that the genomes of present-day human populations outside Africa originate almost entirely from a single major migration out around 50,000-70,000 years ago, accompanied or followed soon after by mixture with Neanderthals contributing ∼2% to the genome of all non-Africans^4,10^. This mixture event is reliably dated from the length of the Neanderthal segments to 7,000-13,000 years before the time when the ancient-DNA-yielding fossil Ust’-Ishim lived (45,000 years ago)^3^. Thus Neanderthal mixture took place 52,000-58,000 years ago, and the migration out of Africa was probably within this time interval or shortly before, so most likely 52,000-60,000 years ago. The admixed population then expanded rapidly over most of Eurasia and Australia^1,2^. As a result, people were present over much of this vast region by 50,000 years ago. The details of this initial expansion, however, remain poorly characterised. Did it follow a coastal route, an inland route, or multiple routes? Where and when did the ancestors of present-day populations begin to diverge? To what extent do present-day populations retain the genetic imprint of these early patterns? Ancient DNA studies using samples 50,000-60,000 years old could potentially provide definitive answers to these questions, but have not so far been reported because of the absence of suitable samples. Genome-wide analyses of present-day populations show a steady decrease in genetic variation with travelling distance from Africa, and have been interpreted in terms of a ‘serial founder’ model which predicts such a decrease^8,9^. While such a pattern may have been initially established in this way, the complexity of subsequent movements and mixing events increasingly documented by ancient DNA from more recent periods^5,11^ suggests that any early pattern of population structure is unlikely to have persisted for >50,000 years. Thus insights from present-day autosomal genomes into the initial out-of-Africa expansion are confounded by the complexity of subsequent prehistory, suitable aDNA is not yet available, and alternative sources of information are needed. The serial founder model nevertheless provides a standard model with which alternatives can be compared.

There is, however, one region of the genome with the potential to inform about these events in depth: the Y chromosome. This is because its male-specific portion provides haplotypes from which a detailed calibrated phylogenetic tree can be created^6^. Several such trees have been constructed independently and are all consistent in being dominated by a massive expansion of non-African Y lineages during the key interval of 50,000-60,000 years ago starting from a single haplogroup designated CT^12-15^ (see Figure 1 for haplogroup designations). Taking into account a rare African D0 lineage and the timeframe summarised above, we have argued^7^ that the initial splits within CT are likely to have occurred in Africa before the exit, and that three lineages, C, D and FT, were carried out by the ancestors of present-day non-Africans. Each of these three lineages subsequently expanded: C and D moderately, and FT massively. We therefore set out to re-examine the early divergences within these three lineages in order to investigate the insights they can provide into male history and perhaps human history more generally in this early period. We assembled available sequences of C, D and FT lineages from worldwide surveys ensuring that common lineages were represented^1,13,14,16,17^, and supplemented them with additional sequences from known rare lineages potentially relevant to early divergences: specifically, Australian C^1,18^, West African D0^7^, Andamanese D^19^, and F chromosomes from China^1^, Vietnam^14^ and Singapore^20^: 1204 sequences in all. We then focussed on the phylogenetic structure of the early divergences within these three lineages, and their geographical distributions revealed by ancient DNA and present-day analyses.

**Figure 1.**
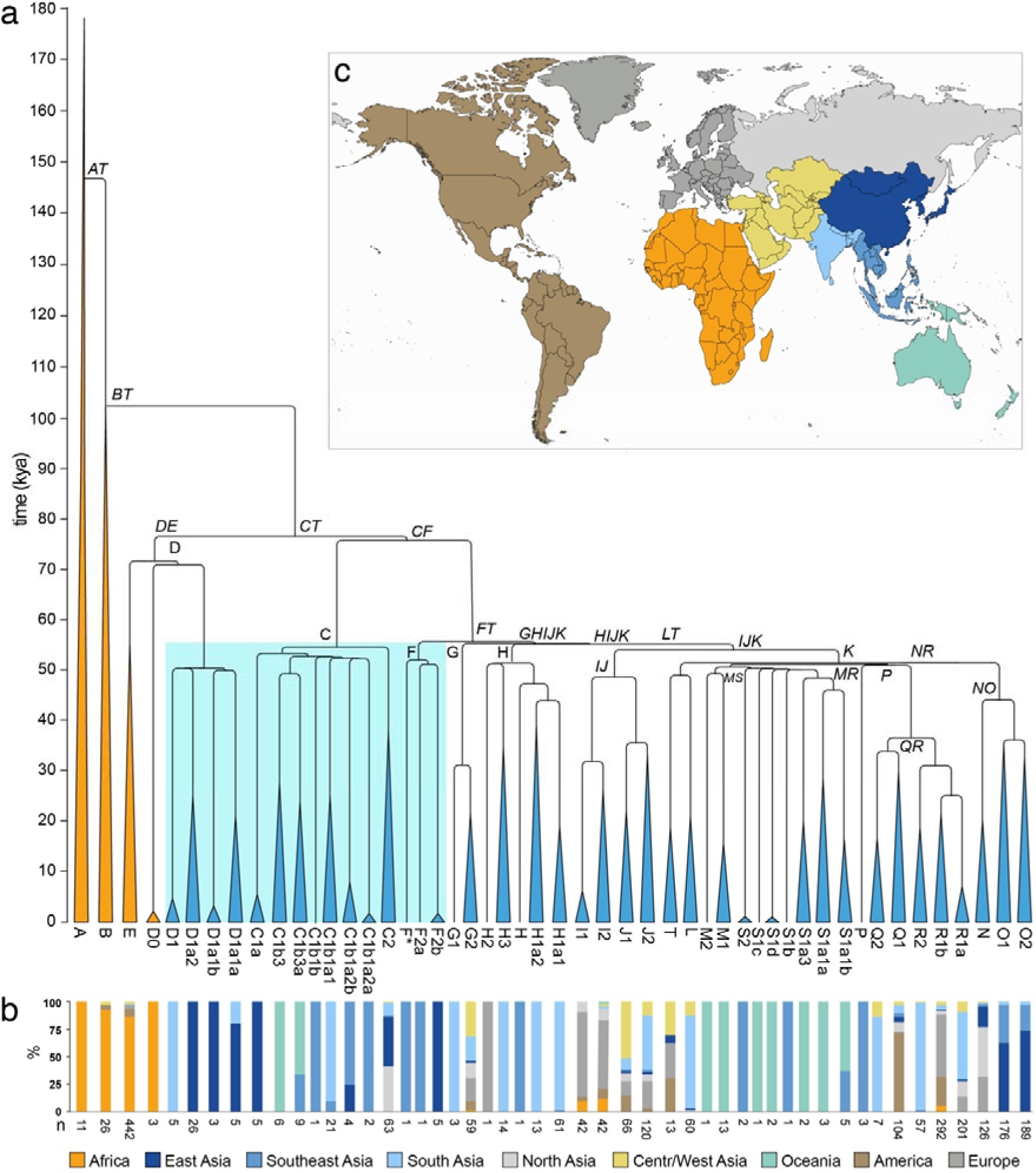
Y-chromosome phylogeny and haplogroup distribution. **(a)** Maximum likelihood Y-phylogeny based on 1204 samples with branch lengths drawn proportional to the estimated times between successive splits according to BEAST analysis. Y lineages currently located in Africa are coloured gold, the others blue. The key lineages of D, C and F are highlighted with a blue box. Haplogroup names indicated in italics correspond to dated splits in Supplementary Table 3. **(b)** Proportion of samples carrying Y lineages shown in (a) coloured according to geographic origin using a total of 2319 samples (1204 samples used to reconstruct the phylogeny plus 1070 non-overlapping samples from the 1000 Genomes Project^14^ and 45 samples from The Singapore Sequencing Malay Project^20^) **(c)** Map showing the geographic divisions used.

The resulting Y-chromosomal tree (Figure 1) depicts 50 lineages, with the African lineages (gold) represented only by the four major African haplogroups without including their subsequent branches, but with the non-African lineages represented more fully to include all those originating before 45,000 years ago and found in the sample of present-day Y chromosomes examined, together with some of the more abundant recent lineages. As expected from previous analyses, this phylogeny shows that the three initial lineages C, D and FT each underwent initial rapid expansions soon after 54,000 years, so that by 50,000 years ago there were seven branches within C, one within D and 15 within FT (23 non-African lineages in all); by 45,000 years ago the number of branches within FT had increased to 24 (36 in all). The branching patterns, together with the present-day locations of the lineages derived from an analysis of 2319 sequences, provide insights into possible locations of the early expansions. Lineage C split into two, C1 and C2; C1 lineages are found today only in East, Southeast and South Asia plus Oceania, while C2 lineages are more widespread and are now found in East and South Asia and also North and Central/West Asia (Figure 2, Extended Data Figure 1). D lineages are entirely confined to East and South Asia. FT lineages now have a worldwide distribution, but the earliest split was into F and GHIJK; F is known only from East and Southeast Asia (Extended Data Figure 1), while GHIJK and its descendants are worldwide. These descendant lineages themselves often have more continent-specific distributions, but 11/12 GHIJK lineages originating before 50,000 years ago have distributions that include East, Southeast or South Asia, apart from a few that are specific to Oceania (Figure 1). Only one (H2, represented by a single sample) is specific to Europe, and none to the region adjoining the likely exit routes from Africa, Central/West Asia, where less than half are now present in the samples examined.

**Figure 2.**
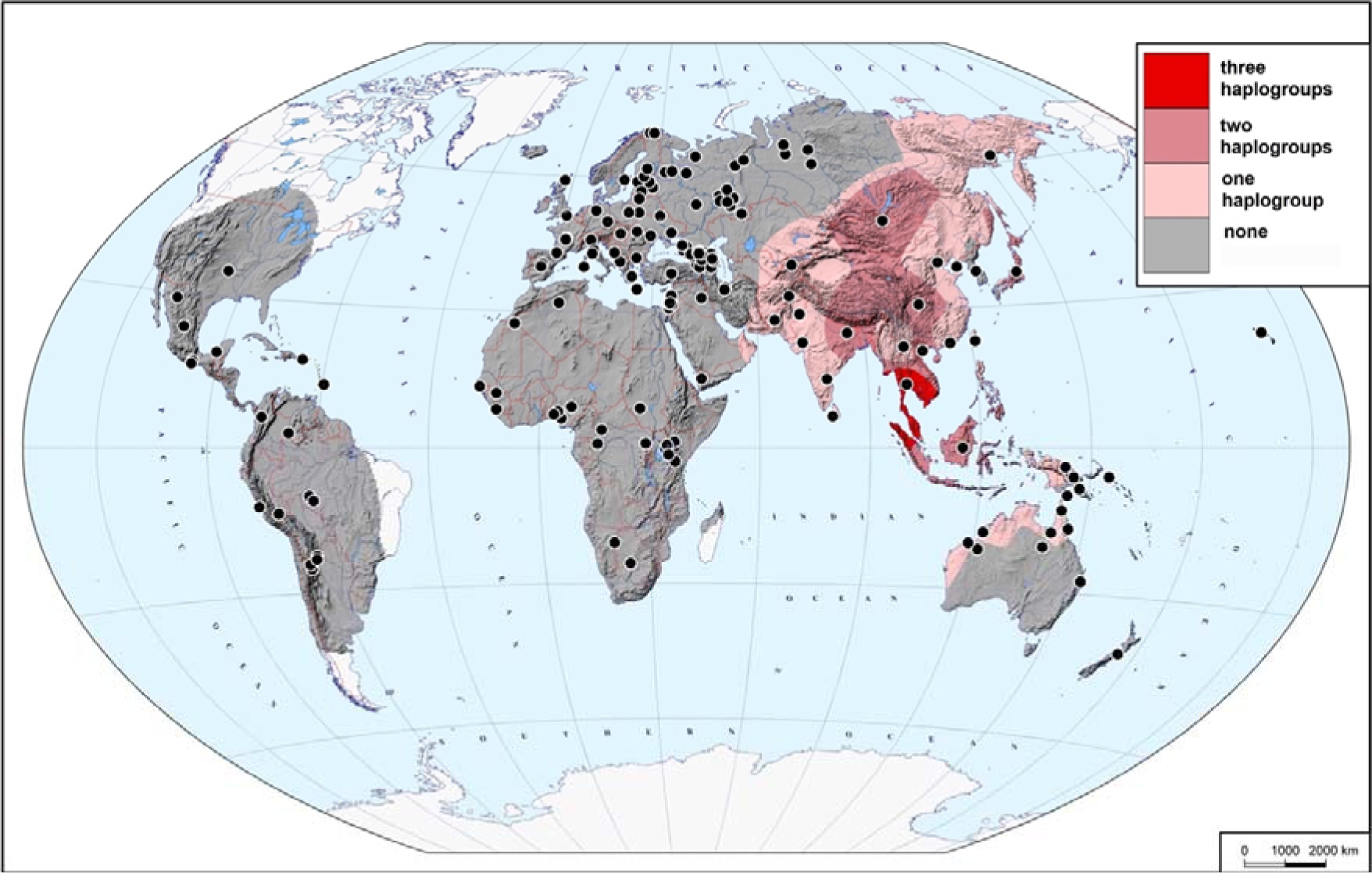
Presence of haplogroups C, D and F in 2302 present-day samples. The map demonstrates how many of the three haplogroups of interest (none, one, two, or all three) were found in different areas of the Old World and Near Oceania. Black dots indicate the locations of the studied populations.

No ancient Y-chromosomal data earlier than 45,000 years ago have been reported, but 21 Asian or European males living 30,000-45,000 years ago are documented, and for 18 of them assignments to C, D or FT have been reported (Extended Data Figure 1, 2, Supplementary table 1)^3,21-26^. Ten belong to the C lineage, six from North Asia and four from Europe. The remaining eight belong to FT, three from North Asia, one from East Asia and four from Europe. Although the data are limited, two conclusions can be drawn. First, none of the ancient samples carry Y lineages outside the 23 represented in Figure 1 at 50,000 years ago. Second, C lineages (both C1a and C1b), now confined to East, Southeast and South Asia plus Oceania, were more widespread 30,000-40,000 years ago, including in Europe where they persisted until after 8,000 years ago^27^, although have now been replaced in Europe by other lineages.

In a simple model of gradual human expansion from Africa to Asia and Oceania without subsequent continental-scale reshaping, we would expect the initial divergences in the Y-chromosomal phylogeny to have occurred in geographical locations close to Africa, and the present-day Y-chromosomal phylogeography to reflect this history by showing the presence of the early-diverging lineages within C, D and FT now being located geographically in Central/West Asia (Figure 3a). In stark contrast, the observed distributions of these lineages all lie further to the East, suggesting that their early divergences occurred in the East (Figure 3b, Extended Data Figure 3), a discrepancy we discuss further below.

**Figure 3.**
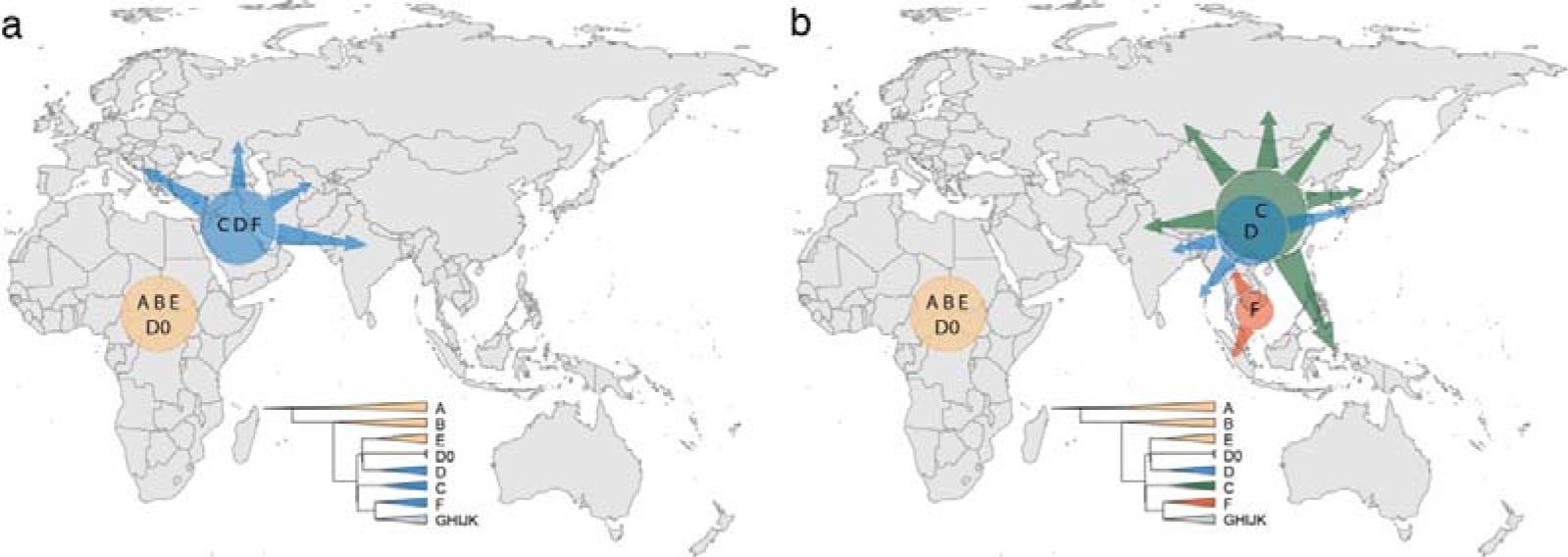
According to serial founder model, the earliest-branching non-African lineages are expected to expand and be present closer to Africa **(a)**, but instead have expanded in East or South-east Asia **(b)**. Simplified Y tree is shown as reference for colours.

The phylogeny of maternally-inherited mitochondrial DNA (mtDNA), like that of the Y chromosome, also retains information from 50,000-60,000 years ago, although with less detail because of its shorter length. Nevertheless, it provides a useful comparison. Outside Africa, the initial split inferred from a combination of ancient and present-day sequences was between lineages M and pre-N, with divergence within M dated to 44,000-55,000 years ago and within N to 47,000-55,000 years ago^28^. Present-day geographical distributions of mtDNAs are less specific than Y chromosomes, and both of these major lineages are widespread worldwide, except that M is absent from present-day Europeans. Nevertheless, M was present in early Europeans until at least 28,000 years ago; moreover, the first branch within the pre-N/N lineage is between the pre-N mtDNA carried by Oase1 from Romania dating to 40,000 years ago (who, incidentally, showed increased Neanderthal admixture) and the remaining worldwide N mtDNAs. mtDNA thus shares with the Y chromosome a history of continental-scale change (loss of M from Europe), in the case of mtDNA dated to after 28,000 years ago. In addition, mtDNA N demonstrates the phylogeographic pattern expected from a simple expansion model, with its earliest divergence in the west.

How then can the present-day Y-chromosomal phylogeography be reconciled with an out-of-Africa expansion? The simplest explanation is that initial western Y chromosomes have been entirely replaced by lineages from further east (Figure 3), perhaps on more than one occasion. This is supported by the more likely origin of GHIJK in the east (Figure 1). Alternative explanations, where the initial divergences within the Y-chromosomal phylogeny did indeed occur in the west, would require either that C, D and F lineages all migrated east, together with some GHIJK lineages, leaving only GHIJK lineages in the west, or that C, D and F were lost by genetic drift in the west, but not in the east. The first alternative would in turn imply unprecedented levels of male-structured migration, and would be difficult to reconcile with subsequent divergences within GHIJK during the next few thousand years, whereby some of the descendent lineages such as G1, H1 and H3 would also need to have migrated east in a male-structured way. The second alternative seems unlikely because genetic effective population sizes have been lower in East Asia than in Europe^29,30^, so less genetic drift is expected in the west. With the ancient DNA and present-day Y-chromosomal data currently available, replacement from the east is therefore the more plausible explanation.

Ancient DNA studies are beginning to show some of the complexity of human genetic history, including providing evidence for large-scale intercontinental movements in the last 30,000 years or so^3,21-26^. The out-of-Africa model requires major intercontinental movements 40,000-60,000 years ago, as well as later expansion into the Americas. From these perspectives, it is perhaps more likely that large-scale movements have continued throughout human prehistory than not. The unique genetic properties of the Y chromosome offer insights into movement during an early period that is currently difficult to investigate in other ways.

## Methods

### Data

Y-chromosomal data from high-coverage whole-genome sequenced samples were combined from the following publicly available or published datasets: the Simons Genome Diversity Project (SGDP)^1^, Polaris (https://github.com/Illumina/Polaris), the Human Genome Diversity Project (HGDP)^16,17^, the Andaman Islands samples^19^, haplogroup D0 samples from Nigeria and additional haplogroup D samples from Tibet^7^, Australian haplogroup C samples^18^ and a haplogroup F* Singapore Malay sample SSM072^20^. Fifty low-coverage whole-genome sequenced samples from the 1000 Genomes Project dataset^14^ were included to represent some of the deep-rooting lineages of haplogroups A, C, F, and H that were not present in other datasets. Additionally, 303 publicly available samples^13^ sequenced at Complete Genomics (CG) were included.

The HGDP, SGDP, Simons, Polaris and Tibetan samples had been mapped to GRCh38, and the Australian, haplogroup D0 and 1000 Genomes Project samples to GRCh37. The reads mapping to the Y chromosome from GRCh37-mapped haplogroup D0, Malay and Andaman Islands samples were extracted using picard (v2.7.2), re-mapped to the GRCh38 using bwa mem (v0.7.17)^31^, followed by duplicate removal using samtools (v1.8).

The genotypes of samples mapped to GRCh38 were jointly called using bcftools (v1.8) with minimum base quality 20, mapping quality 20 and defining ploidy as 1, using the 10.3Mb of chromosome Y sequence previously defined as accessible to short-read sequencing^32^. Similarly, samples mapped to GRCh37 were jointly called using identical parameters. The calls were filtered as follows: removing single nucleotide variants (SNVs) within 5 bp of an indel (SnpGap) and removing indels. The genotypes of high-coverage samples with an overall mean read depth on chromosome Y ≥ 12× were filtered for minimum read depth of 3, samples with lower mean read depth for minimum read depth of 2, except that no minimum read depth filter was applied to the 1000 Genomes Project low-coverage samples. Additionally, if multiple alleles were supported by reads, then the fraction of reads supporting the called allele should be ≥0.85; otherwise the genotype was converted to missing data. The CG dataset was obtained as a GRCh37 all-sites vcf file, where all genotypes with the CG-specific VQLOW quality tag had been converted to missing data. All GRCh37-based vcf files were then merged using bcftools, lifted over to GRCh38 using picard followed by merging with the rest of GRCh38-based data. High-coverage samples with ≥5% of missing data across all sites and sites with ≥3% of missing calls across samples were removed using vcftools (v0.1.14). Two samples (CongPy6 and ISR07) from the CG dataset were later removed due to unusually long terminal branches. After filtering, a total of 9,777,468 sites remained, including 82,019 variant sites (47,367 singletons) (Supplementary Dataset 1).

The final dataset includes 1208 samples: 610 from the HGDP, 95 from the SGDP, 126 from the Polaris dataset, 13 Australian aboriginal samples, five samples from the Andaman Islands, 7 haplogroup D samples, 301 CG samples, one Singapore Malay and 50 low-coverage samples from the 1000 Genomes project (Supplementary Table 2).

Two overlapping samples (HG03100 and HG00190) between the Simons and Polaris datasets and two duplicate samples in the CG dataset (Murut5 and Komi2) were retained as internal controls, making it a total of 1204 independent individuals.

In addition, 1070 non-overlapping samples from the 1000 Genomes Project^14^ and 45 samples from The Singapore Sequencing Malay Project^20^ (Supplementary Table 4, 5) were included in the phylogeographic analysis using previously defined Y lineage information.

### Phylogenetic tree construction and dating

The maximum likelihood Y phylogeny including 1208 samples and 82,019 variant sites was inferred using RAxML v8.2.10 with the GTRGAMMA substitution model^33^. The tree was visualized using the FigTree software (v1.4.4) (http://tree.bio.ed.ac.uk/software/figtree/) with midpoint rooting (Extended Data Figure 4).

The ages of the internal nodes in the phylogenetic tree were estimated using both the ρ statistic^34^ and the coalescent-based method implemented in BEAST^35,36^ using only the high-coverage genomes (Supplementary Table 2).

The ρ statistic was estimated as described^18^. Briefly, the pairwise divergence estimates were obtained from the final all-sites vcf, ignoring sites with missing genotypes in either of the samples. If multiple samples were available in a given clade, then per-pair divergence estimates were averaged across them. The divergence times in units of mutations per site were converted to units of years by applying a point mutation rate of 0.76×10^−9^ mutations per site per year^3^. The 95% confidence intervals of the divergence times were estimated using the uncertainty of the mutation rate (0.67–0.86×10^−9^)^3^. To reduce the computational cost, if either group of descendants of the node to be dated contained more than 100 samples, then 1/3 of randomly selected samples were used to obtain the pairwise divergence estimates.

To reduce the computational cost of running BEAST, a smaller dataset containing 332 samples was used for dating. Samples were selected to represent the major branches in the phylogenetic tree and also all the haplogroup C, D and F samples (Supplementary Table 2). An initial maximum likelihood phylogenetic tree was constructed using RAxML with a set of 50,204 variant sites, then using this as a starting tree for BEAST (v1.8.1). Markov chain Monte Carlo samples were based on 116 million iterations, logging every 1,000 iterations. The first 10% of iterations were discarded as burn-in. Eight independent runs were combined using LogCombiner. A constant-sized coalescent tree prior, the HKY substitution model, accounting for site heterogeneity (gamma) and a strict clock with a substitution rate of 0.76×10^−9^ (95% confidence interval: 0.67×10^−9^ to 0.86×10^−9^) single nucleotide mutations per bp per year^3^ was used. A prior with a normal distribution based on the 95% confidence interval of the substitution rate was applied. Only the variant sites were used, but the number of invariant sites was defined in the BEAST xml file. A summary tree was produced using TreeAnnotator (v1.8.1) and visualized using the FigTree software (Extended Data Figure 5).

### Y haplogroup nomenclature

The Y haplogroups of each sample were predicted from the all-sites vcf file with the yHaplo software (https://github.com/23andMe/yhaplo) using a version where the marker coordinates in the relevant input files had been replaced to correspond to the GRCh38 assembly^16^. The identified terminal SNV for each sample was used to update the haplogroup name to correspond to the International Society of Genetic Genealogy nomenclature (ISOGG, https://isogg.org, v03.10.19) (Supplementary Table 2). The exceptions were haplogroup C, D and K*/M samples for which the states (ancestral or derived) of all haplogroup-specific markers included in ISOGG v03.10.19 database were checked and the haplogroup name updated according to the most terminal SNV in derived state. Additionally, for haplogroup D0 samples the original nomenclature^7^ was followed (Supplementary Table 2). For the haplogroup F samples the following nomenclature is suggested to correspond to the phylogenetic tree: sample HG02040 to be defined as F*, SSM072 as F2a and the five Lahu samples (HGDP01317, HGDP01318, HGDP01320, HGDP01321 and HGDP01322) as F2b (defined as F2 according to the ISOGG database).

### Cartographic analysis

The combined dataset of 2302 Y chromosomes from 269 populations with geographic coordinates of origin available (Supplementary Table 5) was used to create the distribution maps of three key haplogroups (C, D, and F). As many populations were represented by very few samples, data on neighboring populations were merged to achieve the average sample size of approximately 50 in the areas with non-zero frequencies of the haplogroups of interest (Supplementary Table 5). The GeneGeo software^37,38^ was used with the generalized Shepard’s method, weight function 3 and radius of influence of 2,000 km to create the grids of interpolated values. The frequency distribution maps (not shown) were created as well as the maps demonstrating presence or absence of a haplogroup (Extended Data Figure 3). In each node of the cartographic grid, the values of these three haplogroup presence maps have been summarized, and combined into a single map (Figure 2) indicating how many of the three haplogroups of interest (none, one, two, or all the three) were found in different areas of the Old World.

## Data Availability statement

Final filtered vcf containing variant sites for 1208 samples is available as Supplementary Data file 1.

## Code Availability

Custom perl code to filter for the allelic read ratio is available at https://github.com/pilleh/chrY

## Acknowledgements

We thank all the donors of the samples for making this work possible.

## Author contributions

C.T-S. and Y.X. initiated and led the study. P.H., A.A. and O.B. performed analyses. C.T-S. and P.H. wrote the manuscript with contributions from all other authors. All authors contributed to the final interpretation of data.

## Tables

**Supplementary table 1** - Ancient male samples living more than 30,000 years ago used in this study

**Supplementary table 2** - Sample information for 1208 individuals

**Supplementary table 3** - Split times estimated using the rho statistic and BEAST

**Supplementary table 4** - Additional samples included for analysis of Y lineage distributions

**Supplementary table 5** - Summary of included populations

## Supplementary data

**Supplementary dataset 1** - final variant sites vcf for 1208 samples

**Extended Data Figure 1.**
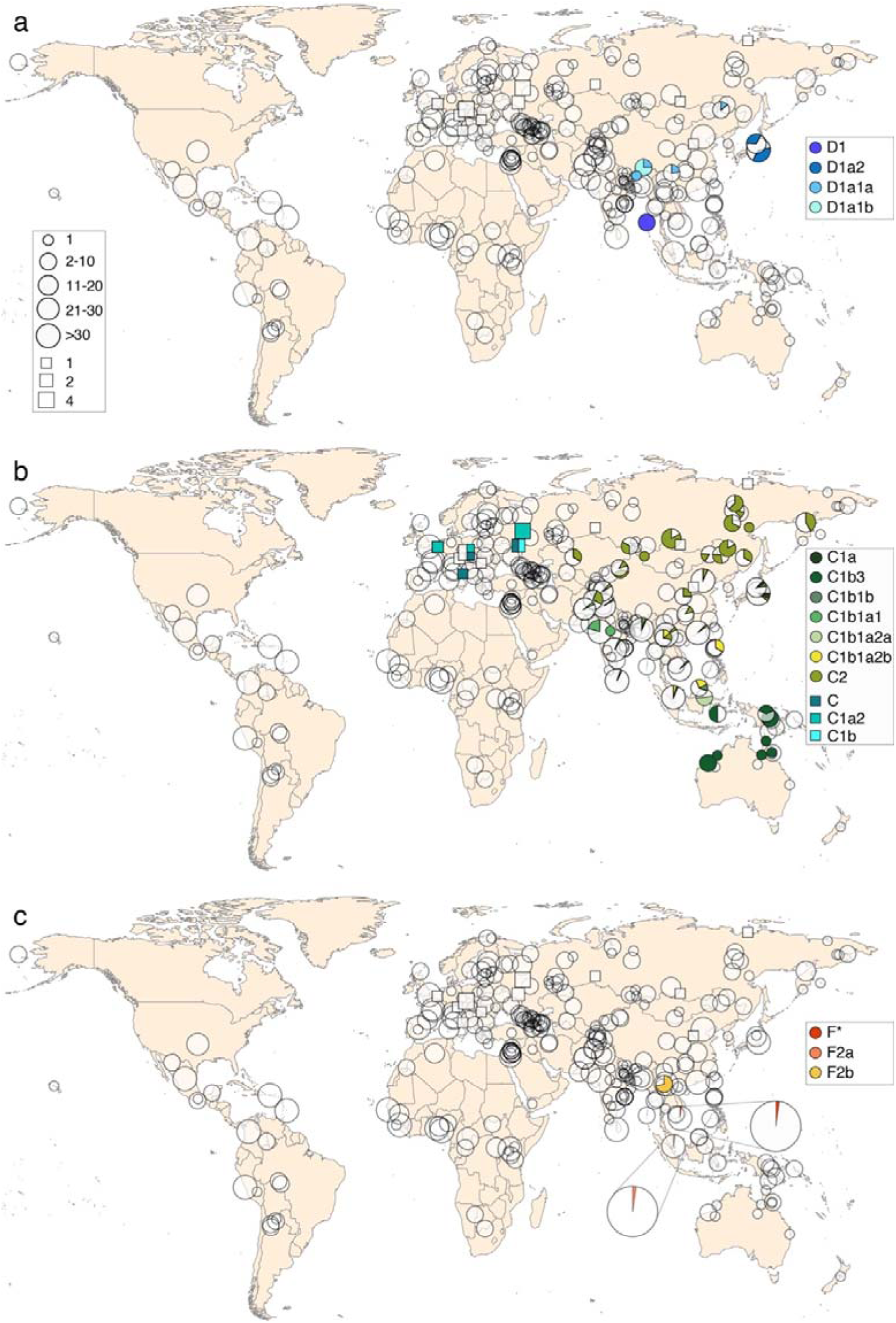
Geographic distribution of males carrying haplogroups D **(a)**, C **(b)** and F **(c).** The geographic origin and approximate sample sizes are shown for 2302 modern samples as circles (for 17 samples from the CG dataset the exact geographic coordinates were not available). Ancient male samples living more than 30,000 years ago are shown as squares, the number of samples corresponding to the size of the square.

**Extended Data Figure 2.**
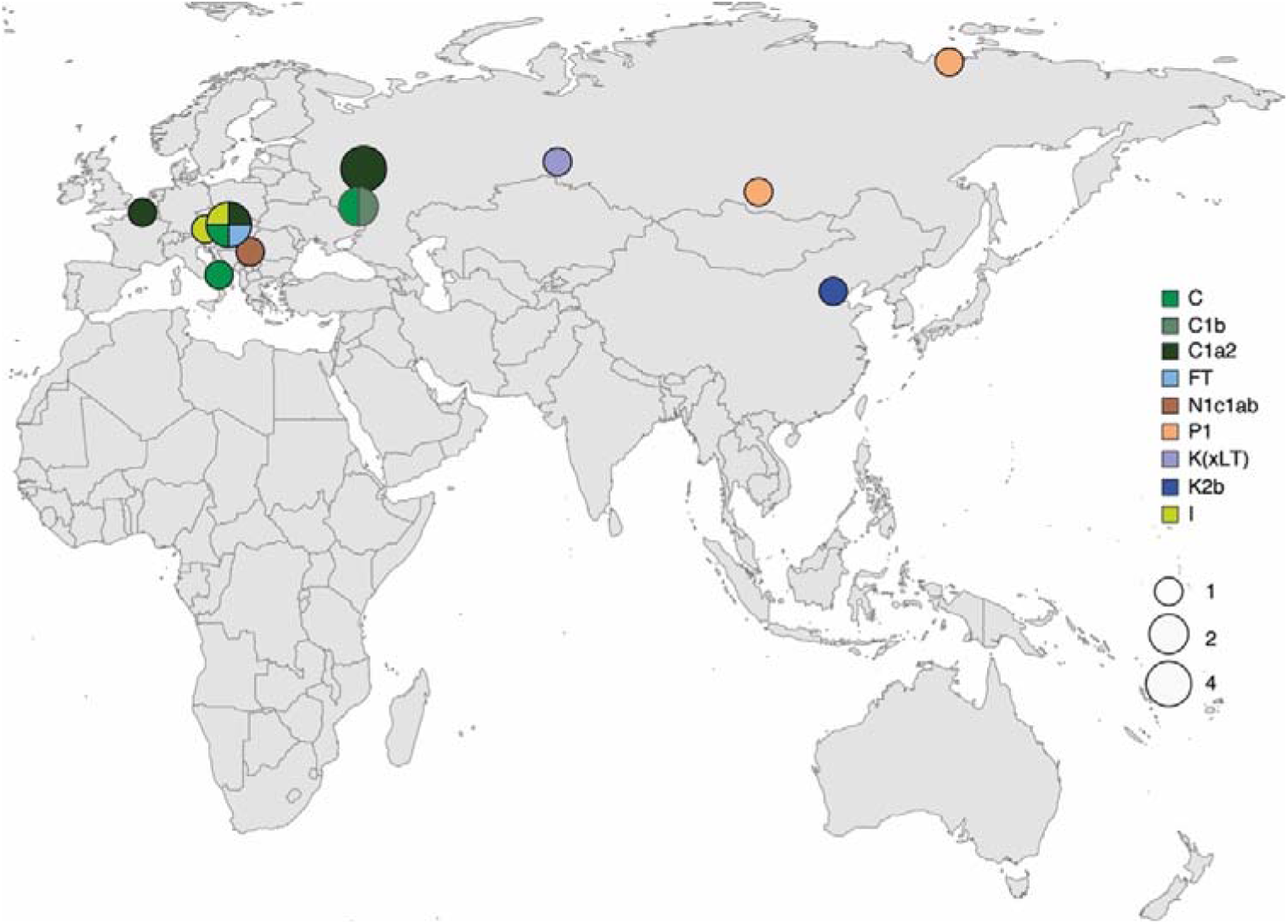
Geographic distribution of Y lineages among ancient male samples living more than 30,000 years ago.

**Extended Data Figure 3.**
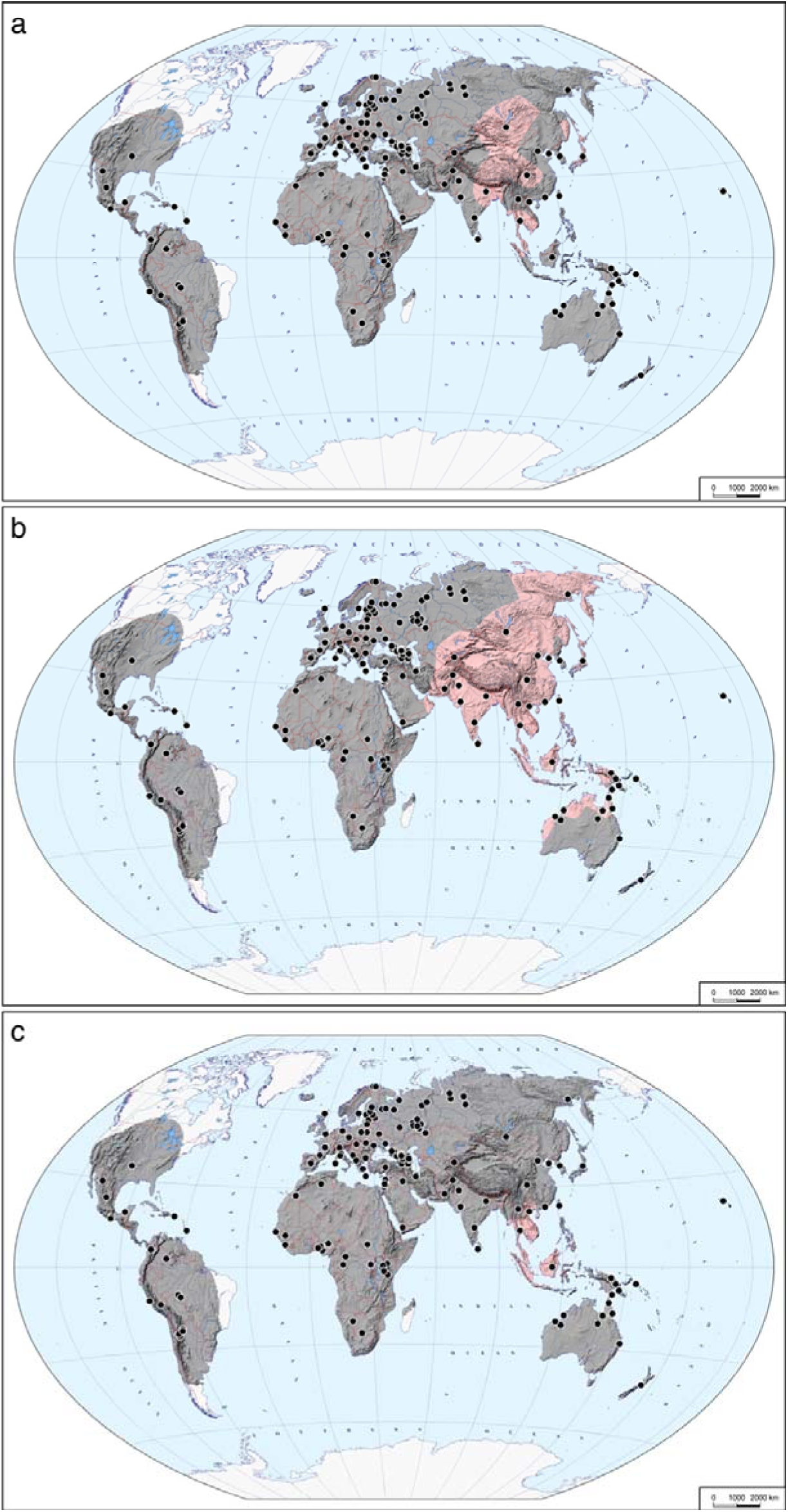
Distribution of haplogroups D **(a)**, C **(b)** and F **(c)** in 2302 samples. Pale red and grey colours indicate the presence or absence of the haplogroup, respectively. Black dots correspond to the geographic origins of the studied populations.

**Extended Data Figure 4.** Maximum likelihood Y-chromosome phylogeny including all 1208 samples used in the study. The sample name in the tree shows the sample ID, population, country of origin and Y haplogroup separated by underscores and is coloured according to the geographic origin of the sample.

**Extended Data Figure 5.** Y chromosome phylogeny based on BEAST analysis of 332 samples. The sample name in the tree shows the sample ID, population, country of origin and Y haplogroup separated by underscores and is coloured according to the geographic origin of the sample. Age and posterior support estimates for main clades are reported in Supplementary Table 3.

