## Supplementary figures and images for "Early replacement of West Eurasian male Y chromosomes from the east"

### SupplementaryFigure4

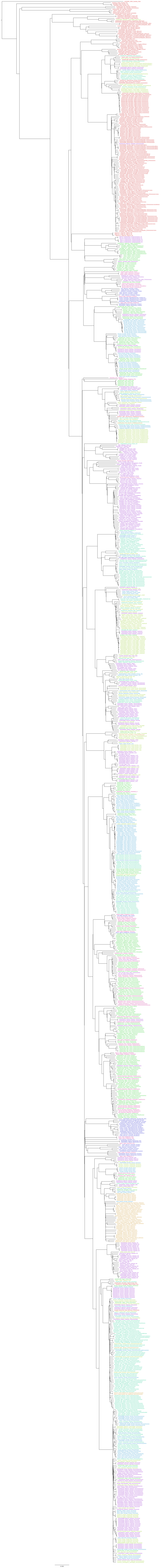

### SupplementaryFigure5

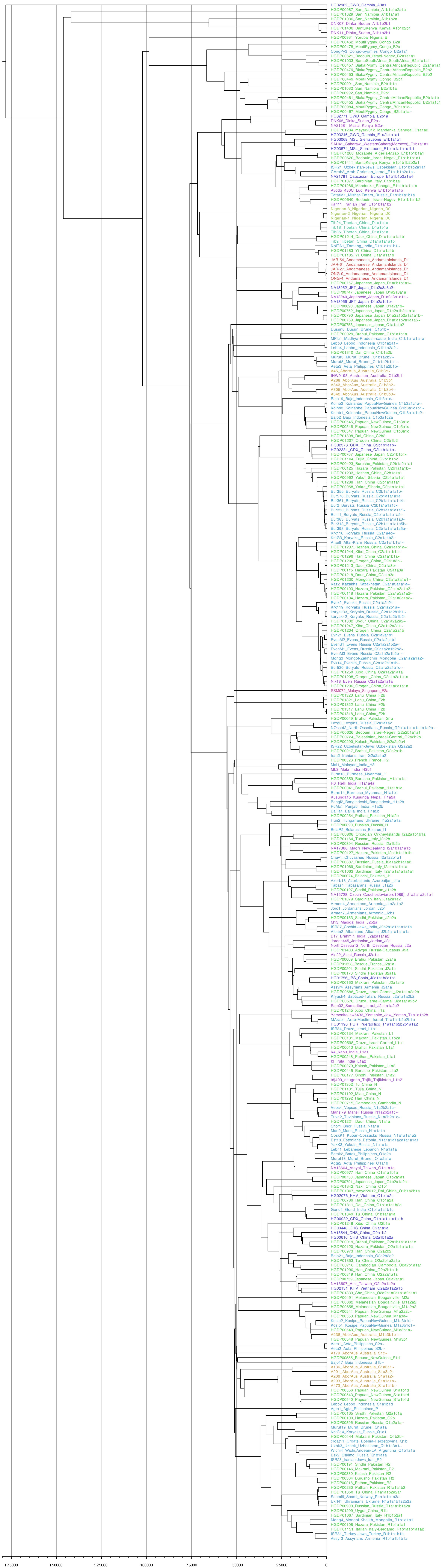
